# Two distinct mechanisms silence *chinmo* in *Drosophila* neuroblasts and neuroepithelial cells to limit their self-renewal

**DOI:** 10.1101/206060

**Authors:** Caroline Dillard, Karine Narbonne-Reveau, Sophie Foppolo, Elodie Lanet, Cédric Maurange

## Abstract

Whether common principles regulate the self-renewing potential of neural stem cells (NSCs) throughout the developing central nervous system is still unclear. In the *Drosophila* ventral nerve cord and central brain, asymmetrically dividing NSCs, called neuroblasts (NBs), progress through a series of sequentially expressed transcription factors that limits self-renewal by silencing a genetic module involving the transcription factor Chinmo. Here, we find that Chinmo also promotes neuroepithelium growth in the optic lobe during early larval stages by boosting symmetric self-renewing divisions while preventing differentiation. Neuroepithelium differentiation in late larvae requires the transcriptional silencing of *chinmo* by ecdysone, the main steroid hormone, therefore allowing coordination of NSC self-renewal with organismal growth. In contrast, *chinmo* silencing in NBs is post-transcriptional and does not require ecdysone. Thus, during *Drosophila* development, humoral cues or tissue-intrinsic temporal specification programs respectively limit self-renewal in different types of neural progenitors through the transcriptional and post-transcriptional regulation of the same transcription factor.

**SUMMARY STATEMENT:** Here, we demonstrate that the transcription factor *chinmo* acts as a master gene of NSC self-renewal in the different regions of the developing *Drosophila* brain where it is controlled by distinct regulatory strategies.

## INTRODUCTION

Limitation of stem cell self-renewal during development ensures that organs reach their appropriate size. However, little is known about the temporal cues and downstream effectors that control stem cell activity during the early steps of tissue building. Recently, the chromatin-associated high mobility group protein HMGA2 has been shown to promote progenitor self-renewing potential in various mammalian tissues during development (Copley et al., 2013; Nishino et al., 2008; Parameswaran et al., 2014). During embryonic and early fetal stages, the RNA-binding proteins (RBPs) Imp1 and Lin28 post-transcriptionally promote *Hmga2* expression in mouse cortical progenitors. In contrast, during late fetal stages, the microRNA let-7 promotes the progressive silencing of *Hmga2* facilitating the termination of self-renewal in the cortex (Nishino et al., 2013; Yang et al., 2015). A similar post-transcriptional mechanism regulates *Hmga2* and self-renewal in fetal hematopoietic progenitors (Copley et al., 2013). In addition, in such progenitors, the transcription factor RUNX1 is also known to silence *Hmga2* during development (Lam et al., 2014). Thus, both transcriptional and post-transcriptional mechanisms operate to regulate the temporal expression of *Hmga2* in the various progenitors allowing limited and controlled self-renewal during development. A strict control of these processes is essential as the deregulation of *Hmga2* can promote unlimited self-renewal and tumorigenesis in these tissues (Fusco and Fedele, 2007). Yet, the mechanisms that regulate the temporal expression of *Lin28*, *Imp1*, *Let-7* or *Runx1* in NSCs and other progenitors during fetal development are still unclear. Moreover, although Hmga2 appears quite widely expressed in the central nervous system (CNS) during early mammalian development, it is still unclear whether the same temporal and regulatory mechanisms operate in the various regions of the CNS to limit NSC self-renewal.

The development of the *Drosophila* CNS is simpler than its mammalian counterpart, and is better understood. As such it represents a good model to investigate the basic principles limiting NSC self-renewal (Homem and Knoblich, 2012). Like *Hmga2* in mammalian cortical progenitors, the BTB Zinc Finger gene *chinmo* is highly expressed in *Drosophila* asymmetrically-dividing NSCs of the ventral nerve cord (VNC) and central brain (CB), called neuroblasts (NBs), during early development, and its silencing during late development is necessary to limit NB self-renewing potential (Narbonne-Reveau et al., 2016). Interestingly, *chinmo* is also regulated at the post-transcriptional level in mushroom body neurons by the RNA-binding proteins (RBPs) Imp and Syncrip, and the let-7 miRNA (Kucherenko et al., 2012; Liu et al., 2015; Wu et al., 2012; Zhu et al., 2006). Similarly, Imp and Lin28 promote *chinmo* expression in NB tumors (Narbonne-Reveau et al., 2016). The post-transcriptional regulation of *chinmo* may be a general feature of NBs as they also co-express *Imp* and *lin28* during early larval stages, and *Syncrip* at later stages (Narbonne-Reveau et al., 2016; Syed et al., 2017). In particular, both *chinmo* and *Imp* need to be silenced during development to allow timely termination of NB selfrenewal before adulthood (Narbonne-Reveau et al., 2016). Upstream of *chinmo, Imp* and *lin28* lies a series of sequentially expressed transcription factors, known as temporal transcription factors for their ability to specify the birth-order of the various NB progeny generated upon successive asymmetric divisions (Isshiki et al., 2001; Kambadur et al., 1998). The sequential expression of temporal transcription factors is used as a timing mechanism to schedule, during late larval stages, the end of the Lin28^+^/Imp^+^/Chinmo+ expression window (Maurange et al., 2008; Narbonne-Reveau et al., 2016). Indeed, blocking sequential expression of temporal transcription factors leads to aberrant maintenance of Chinmo, Imp and Lin28, triggering unlimited NB self-renewal in adults (Maurange et al., 2008; Narbonne-Reveau et al., 2016).

Concomitant to NB asymmetric divisions in the VNC and CB during larval stages, a neuroepithelium (NE) first expands and then undergoes differentiation into neural progenitors that will form the optic lobes (OLs) in the brain. Part of this NE, named the outer proliferation center (OPC), will be converted into short-lived NBs that will generate the neurons of the medulla (Egger et al., 2007; Lanet et al., 2013; Yasugi et al., 2008). The NE-to-NB conversion in the OPC is initiated around mid-L3 by high levels of ecdysone produced by the ring gland after the larva reaches a critical weight (Lanet et al., 2013; Lanet and Maurange, 2014). Indeed, in addition to commit larvae to metamorphosis, ecdysone in the brain triggers the rapid progression of a differentiation wave throughout the NE, allowing the rapid differentiation of all NE cells into NBs (Lanet et al., 2013; Yasugi et al., 2010; Yasugi et al., 2008). Inactivation of ecdysone signaling in NE cells leads the unlimited persistence of a proliferative NE in adults. By limiting the self-renewal capacity of NE cells and promoting their rapid differentiation in NBs, ecdysone therefore limits the number of medulla NBs produced, consequently allowing the optic lobe to reach an appropriate final size (Lanet et al., 2013). Moreover, because ecdysone is produced in large quantities once the larvae has reached a critical mass, this mechanism coordinates the initiation of differentiation with organismal growth (Lanet and Maurange, 2014; Layalle et al., 2008; Mirth et al., 2005).

Thus, both cell-intrinsic and systemic signals are used to limit neural progenitors self-renewing potential in the different regions of the developing CNS in *Drosophila*. Yet, it remains unclear whether similar effectors downstream to ecdysone or to the temporal transcription factor series operate in the various types of neural progenitors.

Here we find that Chinmo not only regulates self-renewal in NBs but also in NE cells during early development. However, while the temporal regulation of chinmo in NBs relies on a post-transcriptional mechanism mainly controlled by a cell-intrinsic timer, its regulation in the NE is transcriptional and controlled by ecdysone. This bi-modal regulation of Chinmo allows NSC self-renewal to be promoted by the same master gene but controlled by different temporal strategies in the various regions of the brain.

## RESULTS

### *chinmo* is expressed in both NBs and NE cells but is silenced at different times

While investigating the role of Chinmo in NBs of the VNC and CB, we noticed that it was also expressed and temporally regulated in the medulla NE. We performed a precise time course to investigate the temporal dynamics of *chinmo* expression in NBs relative to the NE. In VNC and CB NBs, *chinmo* is expressed from larval hatching up to the early L3 stage (Fig. 1A-C). However, we find that the silencing of *chinmo* in these NBs is not synchronous, suggesting a NB-intrinsic timing mechanism that is not coordinated between NBs (Fig. 1B). In the NE, *chinmo* is expressed from larval hatching, but remains expressed longer than in most NBs, undergoing a rapid and synchronous silencing around mid-L3 stages (between 12 and 24 hr after the L2/L3 molt, Fig. 1C,D). Chinmo is also expressed for a short period of time in the first few medulla NBs converted from the NE around this time (Fig. 1C, asterisk). Thus, Chinmo in NE cells is silenced synchronously suggesting that a systemic signal may coordinate synchronous Chinmo silencing in this region (Fig. 1E). Together, these observations suggest that the timing of *chinmo* silencing in the NE and NBs is controlled by different temporal mechanisms (Fig.1F).

**Figure 1:**
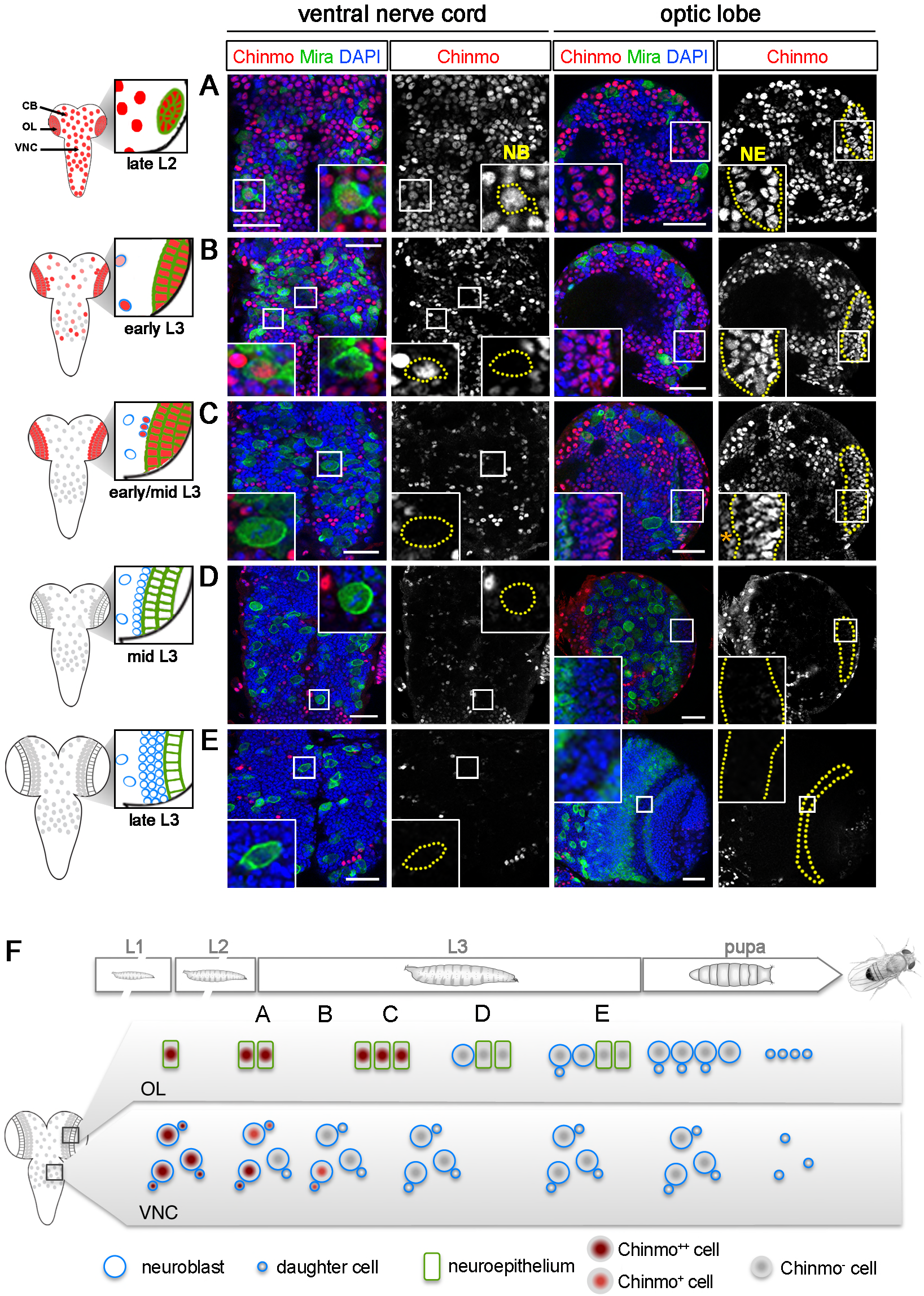
Chinmo is silenced earlier in VNC and CB NBs than in the NE. The scale bar in all images represents 30 μm. NBs are labeled using an anti-Mira antibody (in green), and NE cells are recognized based on their lateral position using the DAPI staining (in blue). Chinmo protein is labeled in red. For each time point, the optic lobes and the ventral nerve cords are pictures from the same CNS. (A) In late L2, Chinmo is highly expressed both in NE cells and in NBs. (B) After the L2/L3 molt, Chinmo progressively and asynchronously disappears in NBs while it is still highly present in the NE. (C) About 10 hr after the L2/L3 molt, most NBs do not exhibit Chinmo whereas it is still observed in the NE and in the first few medulla NBs converted from the NE (asterisk). (D) About 20 hr after the L2/L3 molt, Chinmo is absent from the NE. (E) Chinmo is completely absent from both NBs and NE cells in wandering L3. (F) Schematic drawing representing Chinmo expression during development in VNC and CB NBs and in the NE.

### *chinmo* is post-transcriptionally regulated in VNC and CB NBs

*chinmo* expression is known to be post-transcriptionally silenced in mushroom body NBs and neurons (Liu et al., 2015; Zhu et al., 2006). However, its mode of regulation is unclear in most NBs of the VNC and CB as well as in the NE. We find by fluorescent *in situ* hybridization (FISH) that in late L3, chinmo RNA can be detected in VNC NBs (data not shown) and in CB NBs and their surrounding late-born neurons (Fig. 2B, box 1), while the protein Chinmo is not produced at this stage. Moreover, the use of a lacZ enhancer trap inserted in the chinmo first exon and previously used to assess *chinmo* transcriptional activity (Flaherty et al., 2010; Zhu et al., 2006) indicated consistent lacZ expression in early and late L3 NBs (Fig. 2A,C,D). Together, this shows that *chinmo* is transcriptionally active in NBs throughout larval stages. Thus, a post-transcriptional mechanism operates to silence *chinmo* in most, if not all, late L3 NBs of the VNC and CB. To further investigate this question, we generated transgenic *Drosophila* allowing the conditional expression of a construct in which the *mcherry* coding sequence is flanked by the 5′ and 3′ *UTRs* of *chinmo* (named *UAS-mCherry*^*chinmoUTRs*^) (Fig. 2A). When transcribed in the VNC and CB NBs using *nab-GAL4 UAS-mCherry*^*chinmoUTRs*^, we observed by immuno-staining a strong expression of mCherry in NBs and their progeny up to early/mid-L3 (Fig. 2E). mCherry levels then rapidly decrease and the signal becomes almost undetectable in late L3 NBs (Fig. 2F). This contrasts with a GFP transgene (without the *chinmoUTRs*) that is concomitantly expressed at a constant level in NBs throughout larval stages (Fig. 2E,F). Finally, we misexpressed two different chinmo transgenes: *UAS-chinmo*^*FL*^ that contains the ORF and the UTRs and *UAS-HA-chinmo* that contains only the ORF and lacks the 5′ and 3′ UTRs (Fig. 2A). Consistently, Chinmo is absent in most NBs of late L3 larvae when the *UAS-chinmo*^*FL*^ is expressed using *nab-GAL4* (Fig. 2G). In contrast Chinmo is highly expressed in NBs of late L3 larva when the *UAS-HA-chinmo* is expressed, leading to an amplification of NBs (Fig. 2H). Thus, the silencing of *chinmo* in late NBs of the VNC and CB is mainly mediated by a post-transcriptional mechanism through the UTRs.

**Figure 2:**
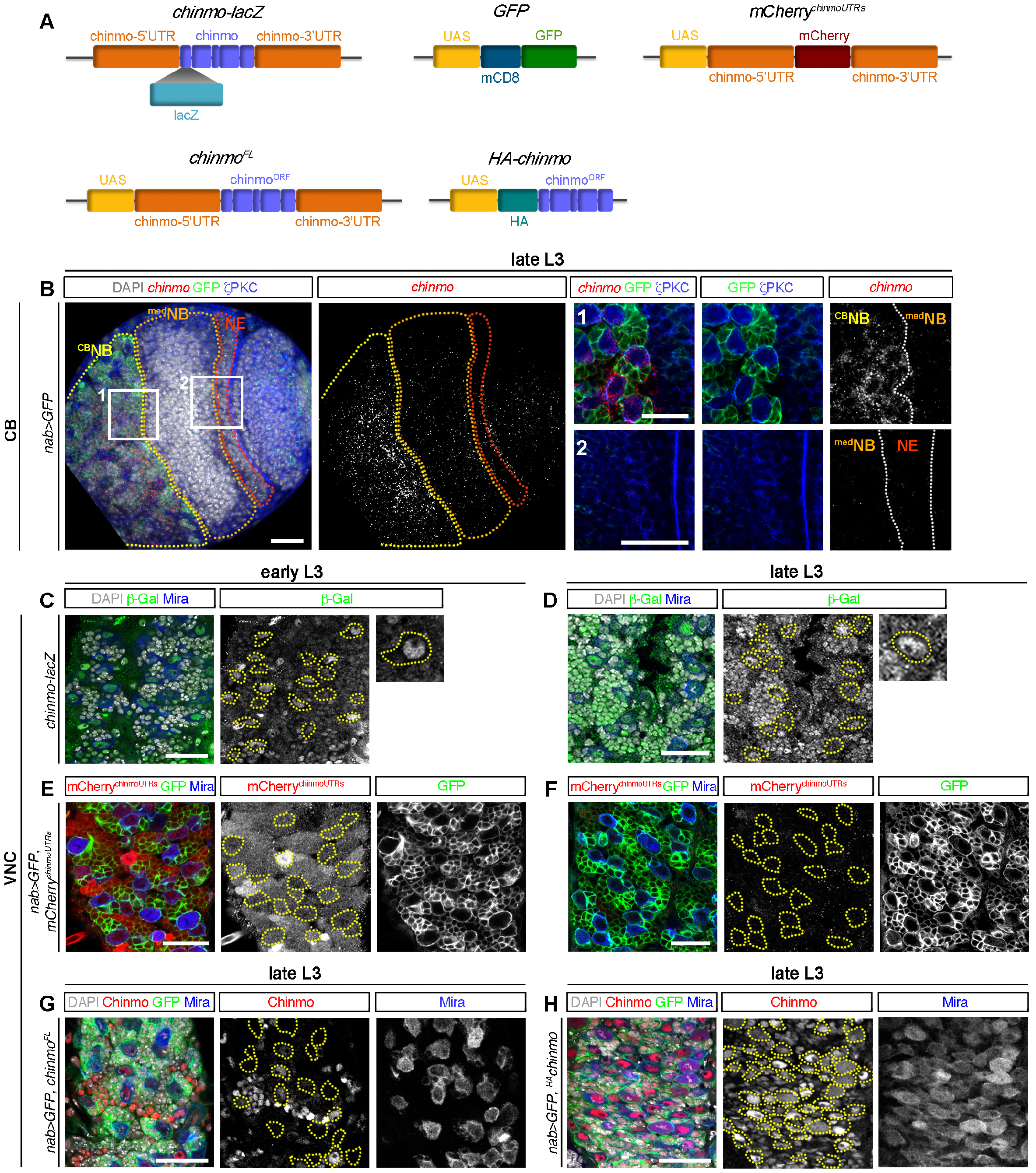
Post-transcriptional regulation of *chinmo* in the VNC and CB NBs. (A) The lacZ insertion into the first exon on chinmo allows the visualization of chinmo transcription. The *mCherry*^*chinmoUTRs*^ transgene recapitulates the post-transcriptional regulation of *chinmo* mediated by its UTRs. The *GFP* transgene serves as a reporter reflecting the transcriptional activity of the GAL4. *Chinmo*^*FL*^ transgene contains the ORF and the UTRs of *chinmo, HA-Chinmo* transgene contains the ORF of *chinmo* only. (B) chinmo transcripts in late L3, as revealed by fluorescent *in situ* hybridization (FISH), exist abundantly in NBs of the CB and their progeny (outlined in yellow, box 1), but are absent from the NE (outlined in red, box 2) and the converted medulla NBs (outlined in orange, box 2). (C-D) Expression of a *chinmo-lacZ* transgene shows that *chinmo* is transcribed throughout development (early L3 -C- and late L3 -D-). NBs are outlined by yellow dotted lines. (E) The *mCherry*^*chinmoUTRs*^ transgene driven by *nab-GAL4* leads to strong mCherry staining in early L3 NBs of the VNC. NBs are also strongly labeled with GFP. (F) mCherry is absent from NBs in late L3 stage, while GFP is still expressed. (G) Chinmo is absent in most NBs of late L3 larvae when *chinmo*^*FL*^ is mis-expressed using *nab-GAL4*. NBs are outlined by yellow dotted lines. (H) Chinmo is highly expressed in NBs when *HA-chinmo* is mis-expressed, leading to an amplification of NBs. NBs are outlined by yellow dotted lines.

### *chinmo* is transcriptionally regulated in the developing NE

We then tested if *chinmo* was also regulated by a post-transcriptional mechanism in the NE. In contrast to VNC and CB NBs, chinmo mRNA is not detected by FISH in NE cells and medulla NBs in late L3 larvae (Fig. 2B, box 2). In addition, when assessing expression of the *chinmo-lacZ* transgene, we found down-regulation of LacZ around mid-L3 in the NE, coinciding with the down-regulation of endogenous Chinmo (Fig. 3A,B). Thus, *chinmo* appears to be transcriptionally silenced in late L3 in the NE and medulla NBs. Moreover, when the *mCherry*^*chUTRs*^ transgene was transcribed in the NE using *ogre-GAL4*, we observed a strong mCherry expression at all stages of larval development (Fig.3C,D). mCherry also persisted in the converted NBs from the NE. Thus, in contrast to VNC and CB NBs, post-transcriptional repression of *chinmo* is not operating in NE cells and medulla NBs (Fig. 2D). Finally, both the misexpression of *UAS-chinmo*^*FL*^ and *UAS-HA-chinmo* in the NE using *ogre-GAL4* lead to high levels of Chinmo protein the NE (Fig. 3E,F) showing that there is no post-translational regulation of *chinmo* expression. All together, these data therefore demonstrate that *chinmo* is regulated by distinct mechanisms in two different regions of the brain. It is post-transcriptionally silenced in most, if not all NBs of the VNC and CB, while it is transcriptionally silenced in the medulla NE and NBs.

**Figure 3:**
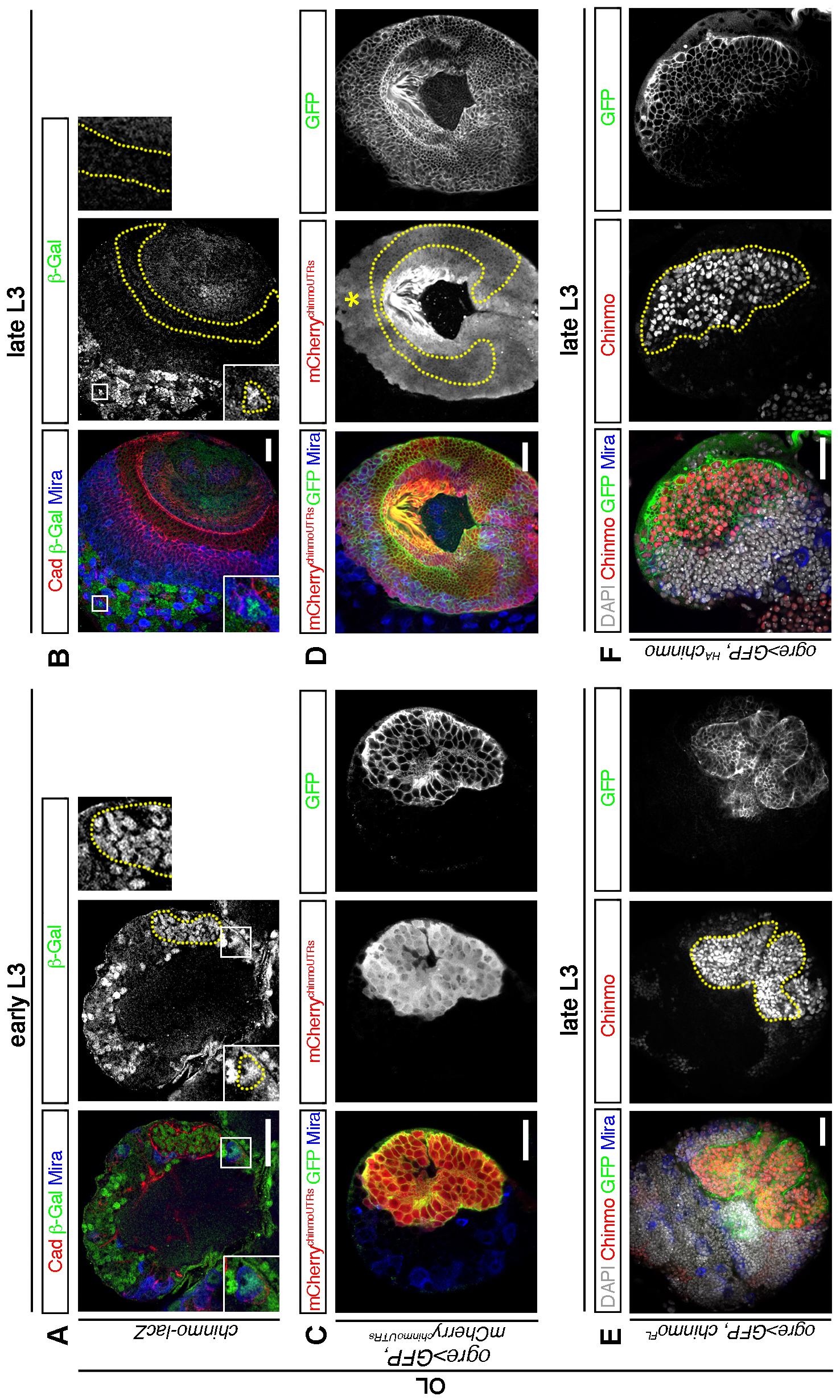
Transcriptional regulation of *chinmo* in the NE. (A-B) *chinmo-lacZ* is active in the NE of early but not late L3 larvae. Note that CB NBs maintain *chinmo – lacZ* expression in late L3 (box). (C-D) mCherry^*chinmoUTRs*^ and GFP driven by *ogre-GAL4* strongly label the NE of early and late L3 The NE is outlined with dotted yellow lines. Note that medulla NBs converted from the NE (asterisk) also express both transgenes. (E-F) The mis-expression of Chinmo^FL^ and HA-Chinmo in the NE using *ogre-GAL4* leads to high levels of Chinmo protein in all epithelial cells.

### Ecdysone signaling is cell-autonomously required to repress *chinmo* transcription in the NE after the CW, but is dispensable in VNC and CB NBs

Because Chinmo down-regulation in the NE coincides with the critical weight (CW) and the subsequent production of ecdysone to initiate metamorphosis, we next tested whether ecdysone signaling could cell-autonomously silence *chinmo*. The ecdysone Receptor (EcR) is continuously expressed in the NE throughout larval stages (Fig. 5E). Strikingly, mis-expression of two dominant-negative forms of EcR (EcR^DN^) (*UAS-EcR.B1-DeltaC655.F645A* and *UAS-EcR.B1-DeltaC655.W650A*) known to efficiently counteract ecdysone signaling (Cherbas et al., 2003), throughout the NE using *ogre-GAL4*, or in clones using MARCM (Lee and Luo, 1999), all led to the maintenance of Chinmo in the NE in late L3 larvae (Fig. 4A-C). Thus, ecdysone signaling cell-autonomously silences *chinmo* around mid-L3 in the NE. In contrast, expression of *EcR*^*DN*^ forms in MARCM and FLP-out clones or using *nab-Gal4* did not lead to the persistence of Chinmo in late L3 NBs, although a slight delay in Chinmo downregulation was observed around early/mid-L3 (Fig. 4E, Fig. S1A,B). Similar results were obtained by abrogating CW-mediated ecdysone pulses using *molting defective* (*mld*^*dts3*^) mutant larvae switched at 29 °C from late L2 (Holden et al., 1986). In that case, we observed persistence of Chinmo in the NE cells of late L3 larvae, but not in VNC and CB NBs (Fig. 4F,G). Thus, in contrast to the NE, ecdysone is not necessary for *chinmo* silencing in late L3 VNC and CB NBs, although it appears to facilitate the timely transition to a chinmo^-^ state (Fig. 4H). These experiments demonstrate that *chinmo* in NBs and in the NE is regulated by different mechanisms. In VNC and CB NBs, it is silenced at the post-transcriptional level by a NB-intrinsic temporal mechanism encoded by the sequential expression of temporal transcription factors. In NE cells, it is silenced at the transcriptional level by mid-L3 pulses of ecdysone produced after the CW.

**Figure 4:**
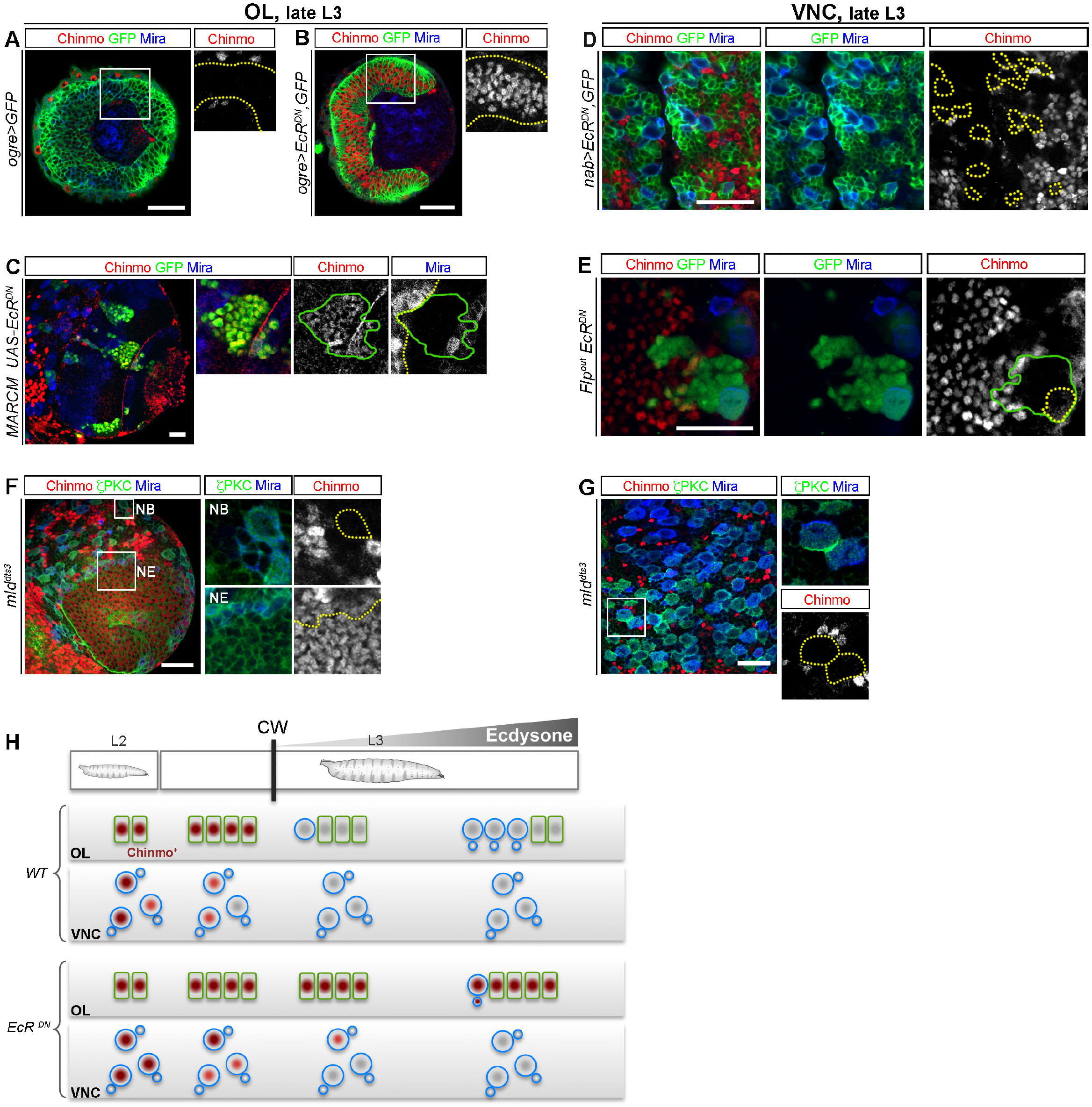
Ecdysone signaling is required for *chinmo* silencing in the NE but is dispensable in VNC and CB NBs. *nab-GAL4 and ogre-GAL4* expressing cells, MARCM clones and Flip-out clones are marked with GFP (green). (A) In late L3, NE cells do not express *chinmo* (red). The NE is outlined in yellow. Medulla NBs are labeled with Mira in blue. Chinmo^+^ cells are glial cells that surround the NE. (B) NE cells mis-expressing *EcR*^*DN*^ fail to silence *chinmo* and to convert into NBs (no Mira^+^ cells). The NE is outlined in yellow. (C) MARCM clones mis-expressing *EcR*^*DN*^ show a delayed NE-to-NB conversion in late L3, as outlined in yellow, and a maintenance of Chinmo. The clone is outlined in green. (D-E) Mis-expression of EcR^DN^ in VNC NBs using *nab-Gal4* (D) or in Flp-out clones (E) does not prevent Chinmo (red) silencing in late L3. NBs are outlined in yellow in D and E and and the clone is outlined in green in E. (F-G) In *mld^dts3^* mutants switch to 29°C in L2 for 48 hr, *chinmo* is silenced in VNC and CB NBs (G), but remains expressed in the NE (F). NBs are labeled with both ζPKC (green) and Mira (blue) whereas NE is labeled with ZPKC only. (H) Schematic recapitulation of the above experiments.

**Figure 5:**
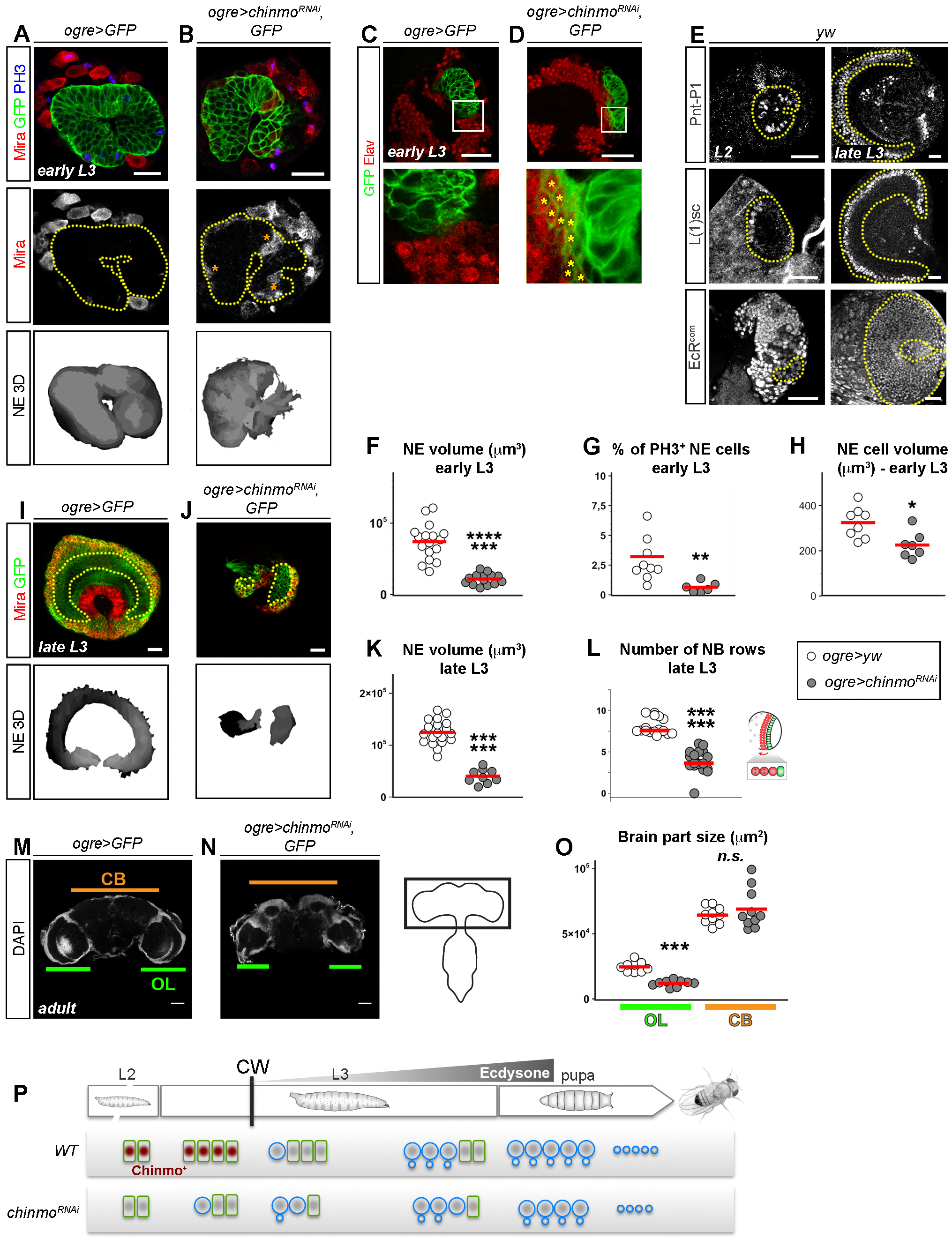
Chinmo promotes NE expansion and protects against precocious differentiation. (A) *ogre>GFP* labels the OPC, that is composed by the NE in early L3. No medulla NBs have been converted. GFP’ Mira^+^ NBs are from the CB. (B) *ogre>chinmo*^*RNAi*^ triggers precocious conversion of NE cells into medulla NBs in early L3 (asterisks), and a reduction of NE size, as highlighted with the 3D reconstruction panels. Mitotic cells are stained with PH3. The yellow dotted line delimits the NE. (C) In early L3, no medulla neurons are present, as observed by the absence of GFP^+^ ELav^+^ neurons. (D) Expression of *ogre>chinmo*^*RNAi*^ leads to the precocious production of GFP^+^ Elav^+^ medulla neurons (asterisks) in early L3, due to the precocious NE-to-NB conversion. (E) The proneural wave markers PntP1 and L(1)sc are expressed in the NE in late L2 before the NB conversion has started. The yellow dotted line delimits the NE. EcR is also expressed in the NE from L2 stages. (F) Volume of the NE in *ogre>GFP* (n=17, m=74004 ± 5857 μm^3^) and *ogre>chinmo*^*RNAi*^, *GFP* (n=14, m=21641 ± 2033 μm^3^) early L3 larvae. *p*-value is 3.02×10^−8^. (G) Mitotic index of NE cells in *ogre>GFP* (n=9, m=3.22 ± 0.56 %) and *ogre>chinmo*^*RNAi*^, GFP (n=6, m=0.64 ± 0.16 %) early L3 larvae. *p*-value is 3.85×10^−3^. (H) NE cell volume in *ogre>GFP* (n=8, m=324 ± 23 μm^3^) and *ogre>chinmo*^*RNAi*^ (n=7, m=± 20 μm^3^) early L3 larvae. *p*-value is 1.28×10^−2^. (I) Late L3 NE (GFP+) and medulla NBs (GFP^+^ Mira^+^). (J) Expression of *chinmo*^*RNAi*^ (ogre>chinmo^*RNAi*^) leads to a small NE (GFP^+^) and fewer medulla NBs (GFP^+^ Mira^+^) in late L3 larvae. The yellow dotted line delimits the NE. (K) Volume of the NE in *ogre>GFP* (n=19, m=124660 ± 5279 μm^3^) and *ogre>chinmo*^*RNAi*^,*GFP* (n=9, m=40196 ± 4362 μm^3^) late L3 larvae. *p*-value is 2.90×10^−7^. (L) Number of converted NB rows along the mediolateral axis (red arrow onthe scheme) in *ogre>GFP* (n=17, m=8.15 ± 0.24) and *ogre>chinmo*^*RNAi*^,*GFP* (n=19, m=3.87 ± 0.30) late L3 larvae. *p*-value is 3.34×10^−7^. (M) Adult control *ogre>yw* flies with optic lobes (OL) and a central brain (CB). (N) Adult *ogre>chinmo*^*RNAi*^ flies have a normal CB but smaller OLs.. (O) OL areas in *ogre>yw* (n=9, m=24621 μm^2^ and SEM=1218 μm^2^) and *ogre>chinmo*^*RNAi*^ (n=9, m=11963 ± 762 μm^2^) adult flies and CB areas in *ogre>yw* (n=10, m=64395 ± 1918 μm^2^) and *ogre>chinmo*^*RNAi*^ flies (n=10, m=69026 ± 4642 μm^2^) adult flies. *p*-value are 4.11×10^−5^ and 0,97, respectively. (P) Schematic recapitulation of the above experiments.

### Chinmo promotes cell growth and counteracts the pre-established proneural front, allowing NE expansion before the CW

We then sought to determine the function of Chinmo in the NE. We noticed that Chinmo down-regulation correlates with the initiation of NE-to-medulla NB conversion triggered by ecdysone after the CW. Thus, shortly after the L2/L3 molt (before the CW), no or rare NE-to-NB conversion is observed (Fig. 5A). In contrast, upon down-regulation of *chinmo* expression by RNAi in the NE from larval hatching using *ogre-GAL4 UAS-chinmo*^*RNAi*^, we observed precocious NE-to-NB conversion in L2 and early L3 (Fig. 5B and Fig. S2A,B,C). This is accompanied by precocious medulla neuron production (Fig. 5C,D). Of note, similar results were obtained using two different RNAi lines provided by TRiP and NIG-Fly, although phenotypes were less penetrant with the NIG-FLY RNAi (Fig. S2B-D). Premature NE-to-NB conversion during early larval stages upon chinmo knock-down could be due to the precocious establishment of the signaling pathways responsible for the pro-neural wave. Alternatively, these pathways may be pre-established during early larval stages and free to operate when chinmo is knocked down from early L2. To investigate this question, we stained the NE for PointedP1 (PntP1) that is downstream of the EGFR pathway and required to initiate and propagate the proneural wave (Yasugi et al., 2010), and for Lethal of Scute (L(1)sc) that labels NE cells at the wavefront (Yasugi et al., 2008). Strikingly, both markers are already expressed in the NE of wild type L2 larvae, demonstrating that the signaling for NE-to-NB differentiation is pre-established early on, before *chinmo* down-regulation (Fig. 5E). Thus, Chinmo in the early NE prevents precocious NE differentiation by blocking the propagation of the pre-established proneural front.

We also found that *chinmo* knock-down led to smaller and less proliferative NE cells in early L3 showing that Chinmo is required for cell growth and to boost mitotic activity (Fig. 5F,G and Fig. S2D). Consequently, down-regulation of *chinmo* in the early NE led to a smaller NE and fewer medulla NBs in late L3 (Fig. 5F, H-L), resulting in a smaller optic lobe in adults (Fig. 5M-O). Thus, together these data show that Chinmo promotes NE expansion before the CW by stimulating cell growth and proliferation and by preventing precocious differentiation (Fig. 5P).

### Temporal regulation of *chinmo* limits the self-renewal of NE cells

We then investigated the impact of a temporal deregulation of *chinmo* expression on the NE. When generating MARCM clones mis-expressing *chinmo* (*chinmo*^*OE*^) from early L2 to late L3 stages, we found a delayed conversion of the NE compared to the surrounding tissue (Fig. 6A). A similar repression of NE conversion was also observed when *chinmo* is mis-expressed in the whole NE (OPC) throughout development (*ogre>chinmo*^*OE*^, Fig. 6B,C,E). However, in such conditions, the *chinmo*^*OE*^ NE was only slightly larger than the *wt* NE in late L3 (Fig. 6B-D). This small difference seemed inconsistent with the strong repression of NE differentiation that is observed (Fig. 6B,C). We detected high levels of apoptosis in *ogre>chinmo*^*OE*^ possibly explaining this phenotype (Fig. 6F). Apoptosis inhibition by mis-expressing Baculovirus *p35* (*ogre>chinmo*^*OE*^, *p35*) led to the massive overgrowth of NE in late L3 with few medulla NBs being converted, consistent with *chinmo’s* ability to prevent differentiation and boost cell growth (Fig. 6G).

**Figure 6:**
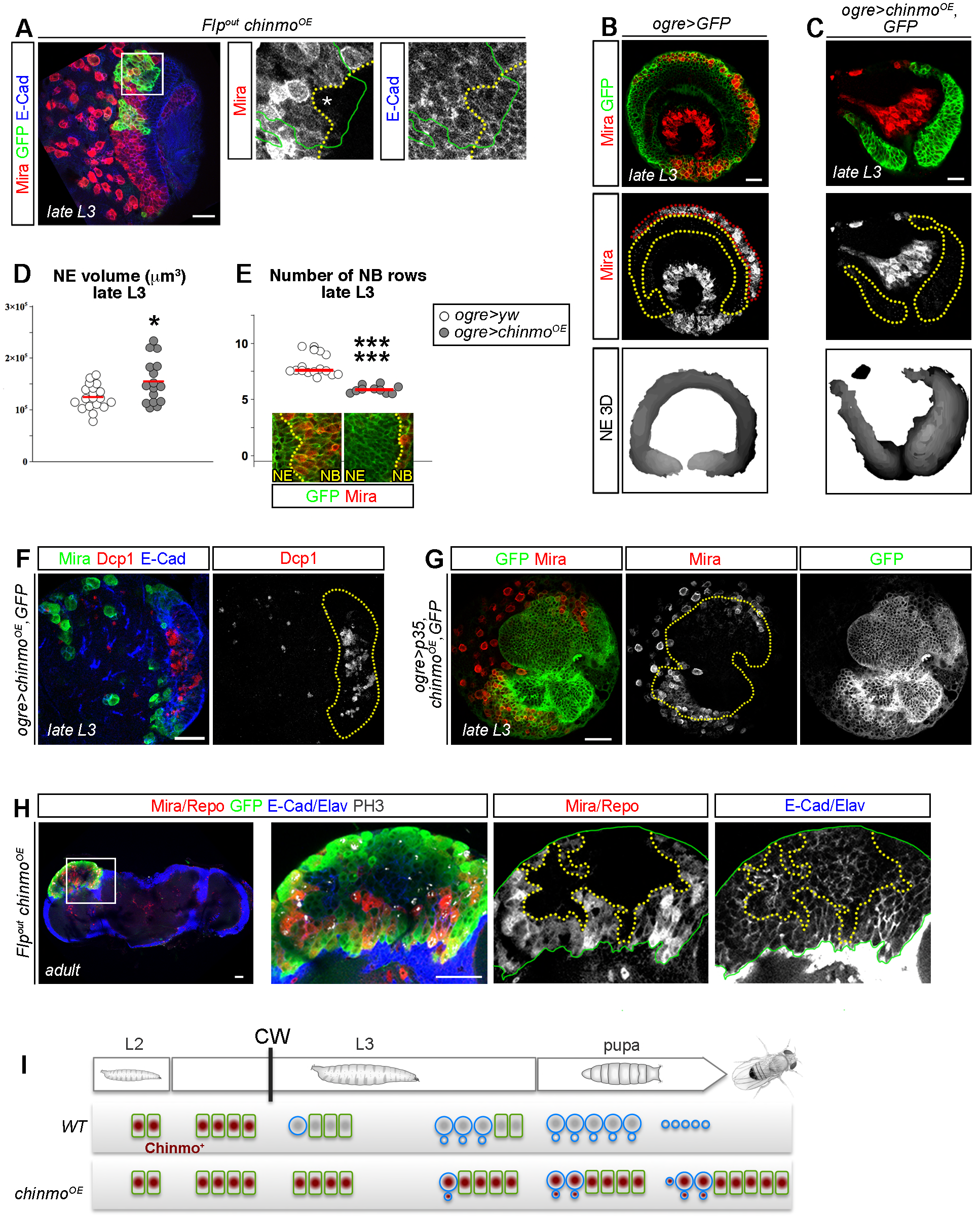
Misexpression of Chinmo delays NE-to-NB conversion and maintains the NE in adults. (A) Misexpression of *chinmo* in *flp-out* clones delays NE-to-NB conversion in late L3 larvae. The clone is delimited by the green line. The pro-neural front is delimited by the yellow dotted line and the delay is marked by the asterisk. (B) Control late L3 larvae. The NE is delineated by the yellow dotted line and converted rows of medulla NBs are highlighted by the red dotted line. (C) Misexpression of *chinmo* in the NE using *ogre-GAL4* prevents conversion of the NE into medulla NBs. (D) Volume of the NE in *ogre>GFP* (n=19, m=124660 ± 5279 μm^3^) and *ogre>chinmo*^*OE*^, *GFP* (n=16, m=154876 ± 10339 μm^3^) late L3 larvae. p-value is 4,06.10^−2^. (E) Number of converted NB rows along the mediolateral axis in *ogre>GFP* (n=17, m=8.15 ± 0.24) and *ogre>chinmo*^*OE*^, *GFP* (n=10, m=5.89 ± 0.10) late L3 larvae. *p*-value is 2,37.10^−7^. (F) *ogre>chinmo*^*OE*^,*GFP* NE cells show high levels of apoptosis, as shown by Dcp1 staining (red). (G) Inhibition of apoptosis by misexpressing *p35* in *ogre>chinmo*^*OE*^ larvae leads to a drastic overgrowth of the NE in late L3 (marked with the GFP), which generates only very few medulla NBs (marked with Mira in red). (H) Misexpression of *chinmo* in *flp – out* clones prevents the total conversion of the NE into NBs and leads to a persisting NE, and NB conversion in the adult OL. (I) Schematic recapitulation of the above experiments.

Strikingly, while the NE is normally eliminated during metamorphosis due to its complete conversion in medulla NBs, mis-expression of *chinmo* in L2-induced MARCM clones prevented the elimination of the NE, leading to perdurance of a proliferative NE in adult optic lobes (Fig. 6H). Thus, down-regulation of Chinmo during development is necessary to allow efficient NE-to-NB conversion leading to NE elimination by the end of development.

### Chinmo does not interfere with the establishment and progression of the temporal transcription factor series in medulla NBs

Medulla NBs converted from the NE sequentially express five temporal transcription factors (Homothorax (Hth), Eyeless (Ey), Sloppy-paired (Slp), Dichaete (D) and Tailless (Tll)) as they age, allowing the generation of a repertoire of neurons (Li et al., 2013). We have observed that the very first medulla NBs generated around the CW transiently express Chinmo (Fig. 7A). We thought to investigate the temporal identity of these NBs and detected Hth in these early-born NBs (Fig. 7A). Thus, Chinmo does not need to be downregulated in medulla NBs to initiate temporal patterning. Moreover, premature NBs induced in early L3 by chinmo knock-down equally initiate and progress throughout the temporal series as we find that they can express Hth, D and Tll (Figure 7B,G,H). In addition, when *chinmo* is misexpressed in the NE, the few medulla NBs that are converted at a low rate appear also able to initiate and progress throughout temporal patterning as they express the temporal factors D and Tll (Fig. 7G,H). It can be noted that because the conversion is delayed when *chinmo* is misexpressed, only very few medulla NBs express the last factor Tll in late L3 compared to the oldest wild-type medulla NBs at the same time (Fig. 7H). All these results show that Chinmo downregulation is not necessary to initiate or to progress through the temporal series in medulla NBs.

**Figure 7:**
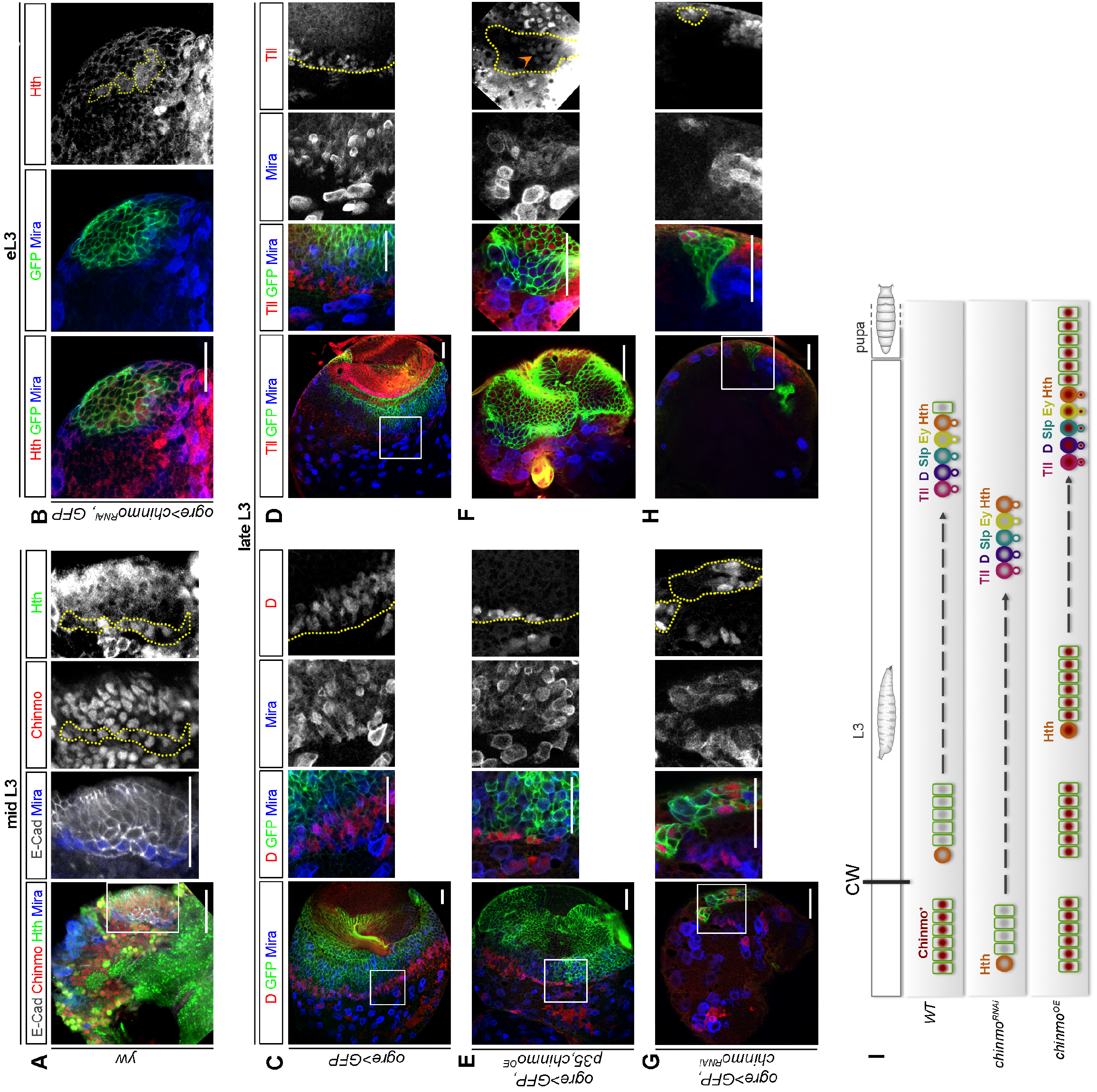
Chinmo does not interferes with temporal series establishment and progression. (A) The first converted medulla NBs in L3 co-express the transcription factors Chinmo (red) and Hth (green). (B) In *ogre>chinmo^RNAi^* brains, premature medulla NBs in early L3 express Hth. (C-H) In late L3, medulla NBs from *ogre>GFP* (C,D), *ogre>chinmo*^*OE*^, *GFP* (E,F) and *ogre>chinmo*^*RNAi*^, *GFP* (G,H) express the late temporal factors D (C,E,G) and Tll (D,F,H). (I) Schematic recapitulation of the above experiments.

## DISCUSSION

Here, we show that self-renewal in the different types of neural progenitors present in the *Drosophila* CNS is promoted during early development by the same transcription factor, Chinmo. However, the expression of chinmo is controlled by different regulatory strategies. This system allows a core self-renewing program to be temporally regulated by distinct intrinsic and extrinsic cues in the various regions of the *Drosophila* brain.

### *chinmo* is a master self-renewal gene in neural progenitors during early *Drosophila* development

We had previously shown that mis-expression of *chinmo* in NBs is sufficient to promote their unlimited self-renewal and that aberrant expression of *chinmo* in NB tumors induced by dedifferentiation is responsible for their unrestricted growth potential (Narbonne-Reveau et al., 2016). These data indicated that Chinmo confers an unlimited self-renewing potential to NBs of the VNC and CB. Silencing of *chinmo* by progression of temporal transcription factors is therefore necessary to limit NB self-renewal during development. Here, we report that *chinmo* is also expressed during early larval stages in the expanding NE of the OPC that will form the medulla region of the optic lobe in the brain. In the NE, Chinmo favors cell growth and proliferation and appears to counteract differentiation. Indeed, loss of Chinmo in the NE of L2 and early L3 larvae is sufficient to induce premature NE-to-NB transition. We have observed that during normal development, L1sc that labels NE cells at the front of the proneural wave, and PntP1, that is downstream to the EGFR pathway and required for the initiation and progression of the pro-neural wave, are already expressed in the NE of L2 larvae, before medulla NBs are being produced. Thus Chinmo appears to counteract the pre-established pro-neural wave in order to prevent precocious NE-to-NB transition, therefore allowing NE expansion from L1 to early L3. The mode of action of Chinmo remains unknown. The knock-down of cell cycle genes has been shown to promote precocious NE-to-NB conversion similar to *chinmo* knock down (Zhou and Luo, 2013). Further work aiming at identifying Chinmo transcriptional targets should help elucidating whether Chinmo prevents differentiation by promoting cell cycle progression and/or by interfering with targets of the EGFR, JAK/STAT, Notch, Hippo signaling pathways that regulate proneural wave progression (Egger et al., 2010; Reddy et al., 2010; Yasugi et al., 2010; Yasugi et al., 2008).

From mid-L3 stages, Chinmo is then silenced. This triggers the sudden acceleration of the pro-neural wave leading to the progressive exhaustion of NE cells through their differentiation into medulla NBs - that generate a much shorter lineage than VNC and CB NBs. In contrast, continuous mis-expression of *chinmo* in the NE induces its continuous expansion throughout larval stages and maintenance of NE self-renewal in the adult brain. Of note, while Chinmo acts as a brake on the NE-to-NB conversion, it does not seem to interfere with the establishment and progression of the series of temporal transcription factors in medulla NBs. Indeed, loss of Chinmo in NE gives rise to precocious medulla NBs that progress through the Hth->Slp->Ey->D->Tll series, similar to the few NBs that can be converted from NE cells mis-expressing *chinmo*. Consistently, the few NBs that are produced and still express *chinmo* in early L3, also exhibit Hth expression.

All together, these data indicate that both in the NE of the OPC and in NBs of the VNC and CB, Chinmo expression confers unlimited self-renewal, and its silencing during larval stages ensures the timely elimination of NBs and NE by the end of development. The general role of a single “master” transcription factor in promoting self-renewal in different types of neural progenitors suggests that the same core set of target genes governs self-renewal, independently of the progenitor type.

Because Chinmo is temporally regulated during development and promotes self-renewal of neural progenitors, it appears to have a role reminiscent to some mammalian oncofetal genes such as *HMGA2*, *MYCN* or *MIZ-1*. These genes all promote neural progenitor proliferation during early development, albeit through different mechanisms. HMGA2 formats chromatin structure, MYCN is a transcription factor that activates genes required for protein biogenesis, and MIZ1 is a Myc cofactor that transforms Myc into a transcriptional repressor of differentiation genes (Boon et al., 2001; Kerosuo and Bronner, 2016; Kishi et al., 2012; Knoepfler et al., 2002; Nishino et al., 2008).

In addition, like Chinmo in VNC and CB NBs, both HMGA2 and MYCN are regulated by RBPs of the IMP (also known as IGF2BP) and LIN28 families in neural progenitors (Bell et al., 2015; Copley et al., 2013; Li et al., 2012; Liu et al., 2015; Molenaar et al., 2012; Narbonne-Reveau et al., 2016; Nishino et al., 2013; Yang et al., 2015). This emphasizes the striking conservation throughout evolution of the post-transcriptional regulation of self-renewal genes by IMP and LIN28 proteins during early development and tumorigenesis.

Therefore, even though no clear ortholog of Chinmo in mammals and of MYCN, HMGA2, and MIZ-1 in insects have been identified (although MIZ1 is a BTB transcription factor with 32% homology with Chinmo), elucidating the mode of action of Chinmo should help to reveal ancestral and generic mechanisms underlying the transcriptional control of stem cell self-renewal.

### *chinmo* is subjected to different modes of temporal regulation in the various regions of the brain

Our work indicates that *chinmo* is under different modes of regulation in NBs and NE cells. In VNC and CB NBs, *chinmo* is post-transcriptionally regulated. Important players in this post-transcriptional regulation could be the RBPs Imp, that could promote *chinmo* expression in early larval NBs, and Syncrip, that could repress *chinmo* in late larval NBs. Both RBPs indeed antagonistically regulate *chinmo* expression in mushroom body neurons and are respectively expressed in early and late NBs (Liu et al., 2015; Syed et al., 2017; Yang et al., 2017). In contrast, *chinmo* is mainly regulated at the transcriptional level in the NE of the optic lobe that expands during early larval development. We had previously shown that ecdysone signaling is strongly activated in the NE shortly after the CW (about 12 hr after the L2/L3 molt) (Lanet et al., 2013). We now demonstrate that one of the main role of ecdysone signaling at this stage is to transcriptionally silence *chinmo*, therefore limiting NE self-renewal and allowing progression of the pro-neural wave. Chinmo down-regulation by ecdysone in the NE does not seem to involve the JAK/STAT pathway, as we did not observe any up-regulation of JAK/STAT activity, by measuring levels of Stat92E (Flaherty et al., 2010), in the EcR^DN^ context (data not shown). In addition, it is likely that ecdysone also regulates other targets in parallel, such as Delta (Lanet et al., 2013) because manipulation of *chinmo* expression alone did not affect Delta expression (data not shown).

Blockage of ecdysone signaling through the mis-expression of different forms of *EcR*^*DN*^, or prevention of ecdysone production in the *mld*^*DTS3*^ context, systematically led to the permanent maintenance of Chinmo in the NE of late L3 larvae. In contrast, in similar conditions, *chinmo* silencing is only delayed by a few hours in VNC and CB NBs. Thus ecdysone is necessary for *chinmo* silencing in the NE but appears dispensable for this purpose in VNC and CB NBs. Instead, in VNC and CB NBs, ecdysone appears to facilitate the timely transition from a Chinmo^+^ to a Chinmo^-^ state that is triggered by temporal series progression (Narbonne-Reveau et al., 2016). Consistent with this is the recent finding that ecdysone promotes some temporal transitions in larval VNC and CB NBs, that in addition to regulate the timely silencing of *chinmo*, are also important for the precise regulation of glial cell numbers in some CB lineages (Syed et al., 2017).

Bi-modal regulation of *chinmo* allows stem cell self-renewal to be under the control of the same master transcription factor, and therefore under the same transcriptional program, while being regulated by different cell-intrinsic or extrinsic cues in the different regions of the brain (Fig. 8). Consequently, self-renewal in the NE appears to be directly controlled by environmental cues such as nutritional conditions and hormones, while other regions of the CNS, like the CB and VNC may be less affected. Interestingly, the regulation of self-renewal by ecdysone signaling in the OL also reveals a mechanism by which endocrine disruptors may affect more heavily the development of specific regions of the brain (Preau et al., 2015).

**Figure 8:**
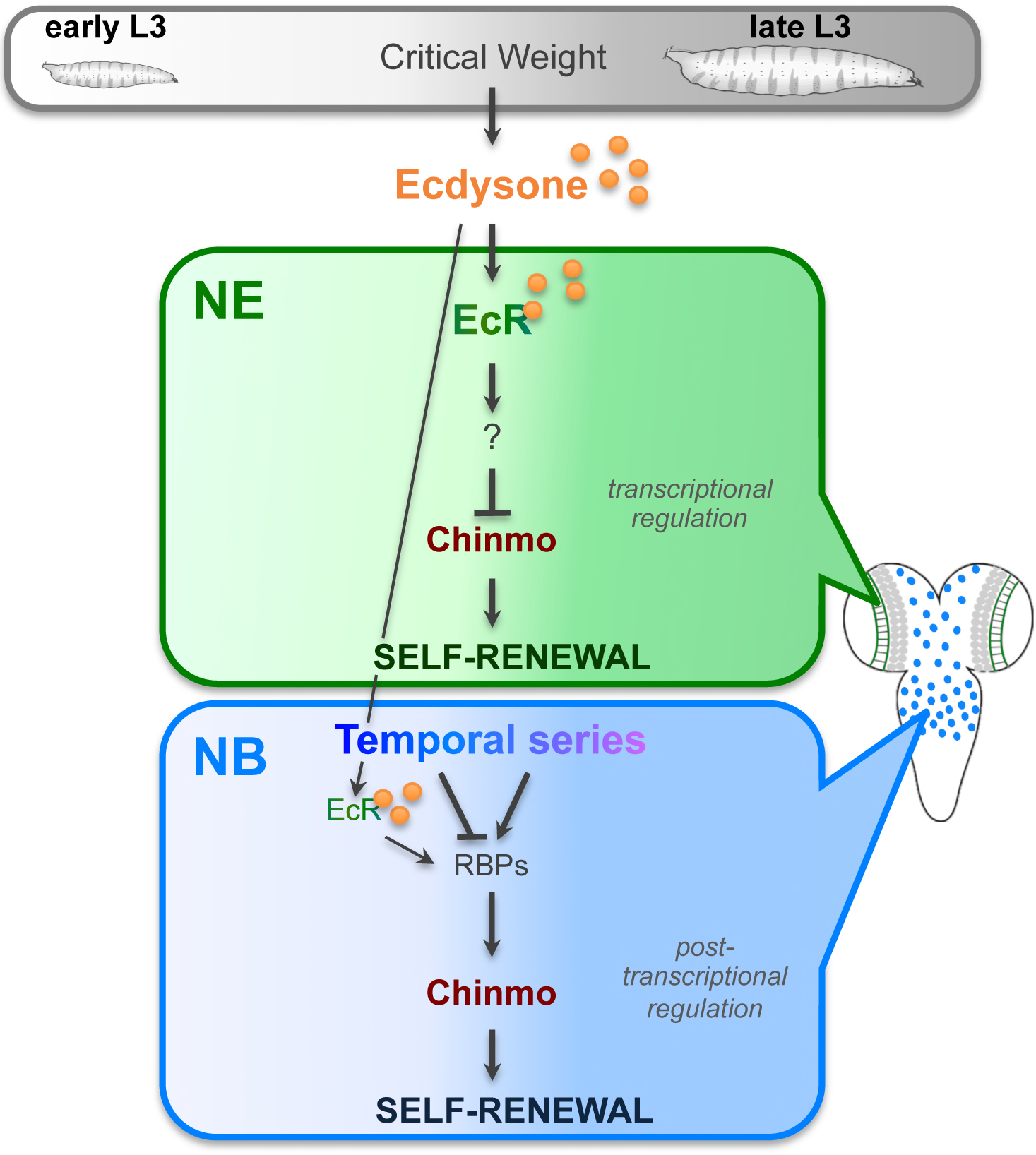
*chinmo* is a master self-renewal gene in neural progenitors during early *Drosophila* development regulated by different mechanisms. Self-renewal of neural progenitors present in the *Drosophila* CNS is promoted during early development by the same transcription factor, Chinmo. However, the expression of *chinmo* is controlled by different regulatory strategies: while *chinmo* during mid-L3 stages is transcriptionally silenced by the Ecdysone pathway in NE cells, its silencing in in the NBs of VNC and CB NBs mainly relies on post-transcriptionnal regulation. Post-transcriptional regulation of *chinmo* in NBs may involve the RBPs Imp, Lin28 and Syncrip that are downstream to temporal transcription factors. Ecdysone signaling facilitates but appears dispensable for *chinmo* silencing in NBs. This system allows a core self-renewing program to be temporally regulated by distinct intrinsic and extrinsic cues in the various regions of the *Drosophila* brain.

Whether the temporal regulation of genes promoting neural progenitor self-renewal during early mammalian development is under the control of a NSC-encoded series of transcription factors or relies on hormonal or more localized cues is still unclear. Interestingly, retinal progenitors in mice have recently been shown to sequentially express orthologs of *Drosophila* temporal transcription factors that specifies the fate of their progeny (Mattar et al., 2015), and NE cells in the developing cortex relies on retinoic acids produced by surrounding meninges (Siegenthaler et al., 2009). Thus, as for *Drosophila*, different temporal mechanisms may govern the self-renewing potential of neural progenitors in the various regions of the developing nervous system in mammals.

## MATERIALS AND METHODS

**Fly culture**. *Drosophila* stocks were maintained at 18 °C on standard medium (8% cornmeal, 8% yeast, 1% agar).

**Image processing.** Confocal images were acquired on a Zeiss LSM780 microscope. FIJI was used to process confocal data, and to compile area and volume data.

**Statistical analysis.** For each experiment, at least 3 biological replicates were performed. Biological replicates are defined as replicates of the same experiment with flies being generated by different parent flies. For all experiments, we performed a Mann-Whitney test for statistical analysis, except for Figure S2D, where a t-test was performed. No data were excluded. Statistical tests were performed with the online BiostaTGV (http://marne.u707.jussieu.fr/biostatgv/). Results are presented as dot plots, also depicting the median in red and a boxplot in the background (Whisker mode: 1.5IQR). The sample size (n), the mean ± the standard error of the mean (m ± SEM), and the *p*-value are reported in the Figure legends.

****: *p*-value ≤ 0.0001,*** : *p*-value < 0.001, **: *p*-value ≤ 0.01 and *: *p*-value ≤ 0.05.

**Fly lines.** Experiments were performed at 29 °C. *yw* line is used as a control. For generating *UAS-EcR.B1-DeltaC655.F645A* (called in this sudy *UAS-EcR*^*DN*^) (Cherbas et al., 2003) MARCM clones (Lee and Luo, 1999), we used *w*, *tub-GAL4*, *UAS-n IsGFP*:: *6xmyc::NLS, hsFLP*^122^; *FRT82B*, *tubP-GAL80/TM6B* crossed to *UAS-EcR*^*DN*^/*CyO_Act_GFP*; *FRT82B/TM6* (from Bloomington #6869). The progeny of the above crosses was heat-shocked 1 hour at 37 °C just after larval hatching and raised at 29 °C. Similar results are obtained with *UAS-EcR.B1-DeltaC655.W650A* (Bloomington #6872). Flip-out clones were generated using *hs-FLP; Act5c>CD2>GAL4, UAS-GFP* (from N. Tapon) with *UAS-chinmo*^*FL*^ (Bloomington #50740) or *UAS-EcR*^*DN*^. The progeny of these crosses was heat-shocked 1 hour at 37 °C just after larval hatching and raised at 29 °C. The GAL4 lines used were the following: *nab-GAL4* (#6190 from Kyoto DGRC, Maurange et al., 2008), *ogre-GAL4* (GMR29C07-GAL4, Bloomington #49340) is active in the OPC, all along larval stages. *wor-GAL4; ase-GAL80* (gift from J. Knoblich) is only active in type II NBs in the central brain. The UAS lines used were: *UAS-chinmo^FL^*, UAS-HA-chinmo (Flaherty et al., 2010), *UAS-EcR*^*DN*^ and *UAS-chinmo*^*RNAi*^ (TRiP #HMS00036, Bloomington #33638 or NIG-Fly #17156R-1). *UAS-dicer2* (Bloomington #24650 and #24651) was used in combination with GAL4 lines in order to improve RNAi efficiency. *UAS-mCD8::GFP* (Bloomington #32186 and #2185) and *UAS-mCD8::RFP* (Bloomington #27399) were used to follow the driver expression. The progeny of the above crosses was raised at 29°C. *chinmo-lacZ* (Bloomington #310440) line was used to monitor *chinmo* transcription. *mld*^*dts3*^ mutant was used to abrogate CW-induced ecdysone pulses (Bloomington #3014).

The larval stages are standardized using morphological criteria. Early L3 are selected just after the L2/L3 molt: the early L3 larvae have the same size than late L2 larvae, but have everted spiracles. Late L3 are selected 48 hours after L2/L3 molt.

**Generation of the *mCherry*^*chinmoUTRs*^ line**. The 5’ and 3’ *Untranslated Transcribed Regions* (*UTRs*) of *chinmo* were cloned on both sides of the *mcherry* coding sequence, under the regulation of *Upstream Activation Sequences* (*UAS*) using the In-Fusion HD Cloning protocol (In-Fusion^®^ HD Cloning kit, Clontech). The entry vector used was pUASTattB-PmeI (a gift from Jean-Marc Philippe, Lecuit lab). The *chinmo 5’UTR* and *3’UTR* sequences were obtained from the EST clone pFLC1-RE59755 (Berkeley Drosophila Genome Project, GOLD collection). The *mcherry* reporter gene was obtained from the pBPGUw-mCherry plasmid (a gift P. Kaspar (Lohmann’s lab))(Sorge et al., 2012). The primers used were:

Ch-5′UTR_F (F1): <u>ATTCGTTAACAGATCT</u>**AGTCAAAAAGAAACTGCCGTG**
Ch-5′UTR_R (R1): <u>GCCCTTGCTCACCAT</u>**GGTGCCAGCAGTGATGCT**
mCherry_F (F2): **ATGGTGAGCAAGGGCGAG**
mCherry_R (R2): <u>TGTTGCGGCTGCTTC</u>**TTACTTGTACAGCTCGTCCATGC**
Ch-3′UTR_F (F3): **GAAGCAGCCGCAACAGCA**
Ch-3′UTR_R (R3): <u>ACAAAGATCCTCTAGA</u>**GGTGAATTTTCATTTGTACGAAGAA**

**Immunohistochemistry**. Dissected tissues were fixed 5 to 15 minutes in 4 % formaldehyde/PBS depending on the primary antibody. Stainings were performed in 0.5 % triton/PBS with antibody incubations separated by several washes. Tissues were then transferred in Vectashield with or without DAPI for image acquisition. Primary antibodies were: chicken anti-GFP (1:1000, Aves #GFP-1020), rabbit anti-RFP (1:500, Rockland #600-401-379), rat anti-RFP (1:500, Chromotek #5F8), mouse anti-Miranda (1:50, A. Gould), rabbit anti-PH3 (1:500, Millipore #06-570), rat anti-PH3 (1:500, Abcam #AB10543), rat anti-Elav (1:50, DSHB #9F8A9), rat anti-DECadherin (1:50, DSHB #DCAD2), mouse anti-Repo (1:200, DSHB #8D12), mouse anti-EcR^com^ (1:7, DSHB #Ag10.2 and #DDA2.7), rabbit anti-βGalactosidase (1:1000, Cappel #559562), rabbit anti-βPKC (1:100, Santa Cruz Biotechnology #sc-216), rat anti-L(1)sc (1:50, A. Carmena), rabbit anti-PntP1 (1:500, J.B. Skeath), rabbit anti-cleaved Dcp-1 (1:500, Cell Signaling #9578), rabbit anti-Tll (1:100, J. Reinitz), guinea-pig anti-D (1:50, (Maurange et al., 2008)), rabbit anti-Hth (1:50, A. Saurin), rat anti-Chinmo (1:500, N. Sokol) and guinea-pig anti-Chinmo (1:500, N. Sokol). Adequate combinations of secondary antibodies (Jackson ImmunoResearch) were used to reveal expression patterns.

**Fluorescent in situ hybridization**. Sens and antisense digoxigenin-labeled riboprobes (DIG RNA Labeling MIX, Roche) against one exonic region of the chinmo transcript (CG31666 RD) were generated (Zhu et al., 2006).

The primers used were:

Chinmo_probe (F1): <u>TAATACGACTCACTATAGG</u>**ACGACCAAGCTGGACAAGAAGCC**
Chinmo _probe (F2): <u>TAATACGACTCACTATAGG</u>**GTTTGGTTTGGTTTGGTTTGGATTTG**

The labeled RNAs were detected by anti-DIG-POD antibody (1:1000, Roche) and visualized with Cy3-tyramide (1:500, PerkinElmer), as previously described (Daul et al., 2010). Tissues were also immunostained using chicken anti-GFP (1:1000, Aves #GFP-1020) and rabbit anti-ζPKC (1:100, Santa Cruz Biotechnology #sc-216) antibodies.

## Acknowledgements

We are grateful to E.A. Bach, A. Carmena, A. Gould, J. Knoblich, T. Lecuit, F. Schnorrer, J.B. Skeath, J. Reinitz, A. Saurin and N. Sokol for flies, plasmids and antibodies. We also acknowledge the Bloomington Drosophila Stock Center (NIH P40OD018537), the Vienna Drosophila RNAi Center (VDRC), TRiP at Harvard Medical School (NIH/NIGMS R01-GM084947), the Berkeley Drosophila Genome Project and the Kyoto DGRC Stock Centers, and the Developmental Studies Hybridoma Bank (DSHB). We thank France-BioImaging/PICsL infrastructure (ANR-10-INSB-04-01). We thank E. Jullian and V. Thomé for technical advices and C. Gaultier and S. Genovese for critical reading of the manuscript. C.D. has been supported by the Ministère de l’Education Nationale, de l’Enseignement Supérieur et de la Recherche; E.L. by the Fondation ARC pour la Recherche sur le Cancer; K.N-R., S.F. and C.M. by the Centre National de la Recherche Scientifique (CNRS).

## REFERENCES

Bell, J. L., Turlapati, R., Liu, T., Schulte, J. H. and Huttelmaier, S. (2015). IGF2BP1 Harbors Prognostic Significance by Gene Gain and Diverse Expression in Neuroblastoma. Journal of clinical oncology: official journal of the American Society of Clinical Oncology 33, 1285–1293.

Boon, K., Caron, H. N., van Asperen, R., Valentijn, L., Hermus, M. C., van Sluis, P.,Roobeek, I., Weis, I., Voute, P. A., Schwab, M., et al. (2001). N-myc enhances the expression of a large set of genes functioning in ribosome biogenesis and protein synthesis. The EMBO journal 20, 1383–1393.

Cherbas, L., Hu, X., Zhimulev, I., Belyaeva, E. and Cherbas, P. (2003). EcR isoforms in Drosophila: testing tissue-specific requirements by targeted blockade and rescue. Development 130, 271–284.

Copley, M. R., Babovic, S., Benz, C., Knapp, D. J., Beer, P. A., Kent, D. G., Wohrer, S., Treloar, D. Q., Day, C., Rowe, K., et al. (2013). The Lin28b-let-7-Hmga2 axis determines the higher self-renewal potential of fetal haematopoietic stem cells. Nat Cell Biol 15, 916–925.

Daul, A. L., Komori, H. and Lee, C. Y. (2010). Multicolor fluorescence RNA in situ hybridization of Drosophila brain tissue. Cold Spring Harb Protoc 2010, pdb prot5462.

Egger, B., Boone, J. Q., Stevens, N. R., Brand, A. H. and Doe, C. Q. (2007). Regulation of spindle orientation and neural stem cell fate in the Drosophila optic lobe. Neural Dev 2, 1.

Egger, B., Gold, K. S. and Brand, A. H. (2010). Notch regulates the switch from symmetric to asymmetric neural stem cell division in the Drosophila optic lobe. Development 137, 2981–2987.

Flaherty, M. S., Salis, P., Evans, C. J., Ekas, L. A., Marouf, A., Zavadil, J., Banerjee, U. and Bach, E. A. (2010). chinmo is a functional effector of the JAK/STAT pathway that regulates eye development, tumor formation, and stem cell self-renewal in Drosophila. Developmental cell 18, 556–568.

Fusco, A. and Fedele, M. (2007). Roles of HMGA proteins in cancer. Nat Rev Cancer 7, 899–910.

Holden, J.J. A., Walker, V. K., Maroy, P., Watson, K. L., White, B. N. and Gausz, J. (1986). Analysis of Molting and Metamorphosis in the Ecdysteroid-Deficient Mutant L(3)3dts of Drosophila-Melanogaster. Dev Genet 6, 153–162.

Homem, C. C. and Knoblich, J. A. (2012). Drosophila neuroblasts: a model for stem cell biology. Development 139, 4297–4310.

Isshiki, T., Pearson, B., Holbrook, S. and Doe, C. Q. (2001). Drosophila neuroblasts sequentially express transcription factors which specify the temporal identity of their neuronal progeny. Cell 106, 511–521.

Kambadur, R., Koizumi, K., Stivers, C., Nagle, J., Poole, S. J. and Odenwald, W. F. (1998). Regulation of POU genes by castor and hunchback establishes layered compartments in the Drosophila CNS. Genes & development 12, 246–260.

Kerosuo, L. and Bronner, M. E. (2016). cMyc Regulates the Size of the Premigratory Neural Crest Stem Cell Pool. Cell reports 17, 2648–2659.

Kishi, Y., Fujii, Y., Hirabayashi, Y. and Gotoh, Y. (2012). HMGA regulates the global chromatin state and neurogenic potential in neocortical precursor cells. Nat Neurosci 15, 1127–1133.

Knoepfler, P. S., Cheng, P. F. and Eisenman, R. N. (2002). N-myc is essential during neurogenesis for the rapid expansion of progenitor cell populations and the inhibition of neuronal differentiation. Genes & development 16, 2699–2712.

Kucherenko, M. M., Barth, J., Fiala, A. and Shcherbata, H. R. (2012). Steroid-induced microRNA let-7 acts as a spatio-temporal code for neuronal cell fate in the developing Drosophila brain. The EMBO journal 31, 4511–4523.

Lam, K., Muselman, A., Du, R., Harada, Y., Scholl, A. G., Yan, M., Matsuura, S., Weng, S., Harada, H. and Zhang, D. E. (2014). Hmga2 is a direct target gene of RUNX1 and regulates expansion of myeloid progenitors in mice. Blood 124, 2203–2212.

Lanet, E., Gould, A. P. and Maurange, C. (2013). Protection of neuronal diversity at the expense of neuronal numbers during nutrient restriction in the Drosophila visual system. Cell reports 3, 587–594.

Lanet, E. and Maurange, C. (2014). Building a brain under nutritional restriction:insights on sparing and plasticity from Drosophila studies. Front Physiol 5, 117.

Layalle, S., Arquier, N. and Leopold, P. (2008). The TOR pathway couples nutrition and developmental timing in Drosophila. Developmental cell 15, 568–577.

Lee, T. and Luo, L. (1999). Mosaic analysis with a repressible cell marker for studies of gene function in neuronal morphogenesis. Neuron 22, 451–461.

Li, X., Erclik, T., Bertet, C., Chen, Z., Voutev, R., Venkatesh, S., Morante, J., Celik, A. and Desplan, C. (2013). Temporal patterning of Drosophila medulla neuroblasts controls neural fates. Nature 498, 456–462.

Li, Z., Gilbert, J. A., Zhang, Y., Zhang, M., Qiu, Q., Ramanujan, K., Shavlakadze, T.,Eash, J. K., Scaramozza, A., Goddeeris, M. M., et al. (2012). An HMGA2-IGF2BP2 axis regulates myoblast proliferation and myogenesis. Developmental cell 23, 1176–1188.

Liu, Z., Yang, C. P., Sugino, K., Fu, C. C., Liu, L. Y., Yao, X., Lee, L. P. and Lee, T. (2015). Opposing intrinsic temporal gradients guide neural stem cell production of varied neuronal fates. Science 350, 317–320.

Mattar, P., Ericson, J., Blackshaw, S. and Cayouette, M. (2015). A conserved regulatory logic controls temporal identity in mouse neural progenitors. Neuron 85, 497–504.

Maurange, C., Cheng, L. and Gould, A. P. (2008). Temporal transcription factors and their targets schedule the end of neural proliferation in Drosophila. Cell 133, 891–902.

Mirth, C., Truman, J. W. and Riddiford, L. M. (2005). The role of the prothoracic gland in determining critical weight for metamorphosis in Drosophila melanogaster. Curr Biol 15, 1796–1807.

Molenaar, J. J., Domingo-Fernandez, R., Ebus, M. E., Lindner, S., Koster, J., Drabek,K., Mestdagh, P., van Sluis, P., Valentijn, L. J., van Nes, J., et al. (2012). LIN28B induces neuroblastoma and enhances MYCN levels via let-7 suppression. Nature genetics 44, 1199–1206.

Narbonne-Reveau, K., Lanet, E., Dillard, C., Foppolo, S., Chen, C. H., Parrinello, H., Rialle, S., Sokol, N. S. and Maurange, C. (2016). Neural stem cell-encoded temporal patterning delineates an early window of malignant susceptibility in Drosophila. Elife 5.

Nishino, J., Kim, I., Chada, K. and Morrison, S. J. (2008). Hmga2 promotes neural stem cell self-renewal in young but not old mice by reducing p16Ink4a and p19Arf Expression. Cell 135, 227–239.

Nishino, J., Kim, S., Zhu, Y., Zhu, H. and Morrison, S. J. (2013). A network of heterochronic genes including Imp1 regulates temporal changes in stem cell properties. Elife 2, e00924.

Parameswaran, S., Xia, X., Hegde, G. and Ahmad, I. (2014). Hmga2 regulates self-renewal of retinal progenitors. Development 141, 4087–4097.

Preau, L., Fini, J. B., Morvan-Dubois, G. and Demeneix, B. (2015). Thyroid hormone signaling during early neurogenesis and its significance as a vulnerable window for endocrine disruption. Bba-Gene Regul Mech 1849, 112–121.

Reddy, B. V., Rauskolb, C. and Irvine, K. D. (2010). Influence of fat-hippo and notch signaling on the proliferation and differentiation of Drosophila optic neuroepithelia. Development 137, 2397–2408.

Siegenthaler, J. A., Ashique, A. M., Zarbalis, K., Patterson, K. P., Hecht, J. H., Kane, M. A., Folias, A. E., Choe, Y., May, S. R., Kume, T., et al. (2009). Retinoic acid from the meninges regulates cortical neuron generation. Cell 139, 597–609.

Sorge, S., Ha, N., Polychronidou, M., Friedrich, J., Bezdan, D., Kaspar, P., Schaefer, M. H., Ossowski, S., Henz, S. R., Mundorf, J., et al. (2012). The cis-regulatory code of Hox function in Drosophila. The EMBO journal 31, 3323–3333.

Syed, M. H., Mark, B. and Doe, C. Q. (2017). Steroid hormone induction of temporal gene expression in Drosophila brain neuroblasts generates neuronal and glial diversity. Elife 6.

Wu, Y. C., Chen, C. H., Mercer, A. and Sokol, N. S. (2012). Let-7-complex microRNAs regulate the temporal identity of Drosophila mushroom body neurons via chinmo. Developmental cell 23, 202–209.

Yang, C. P., Samuels, T. J., Huang, Y., Yang, L., Ish-Horowicz, D., Davis, I. and Lee, T.(2017).Imp and Syp RNA-binding proteins govern decommissioning of Drosophila neural stem cells. Development.

Yang, M., Yang, S. L., Herrlinger, S., Liang, C., Dzieciatkowska, M., Hansen, K. C.,Desai, R., Nagy, A., Niswander, L., Moss, E. G., et al. (2015). Lin28 promotes the proliferative capacity of neural progenitor cells in brain development. Development 142, 1616–1627.

Yasugi, T., Sugie, A., Umetsu, D. and Tabata, T. (2010). Coordinated sequential action of EGFR and Notch signaling pathways regulates proneural wave progression in the Drosophila optic lobe. Development 137, 3193–3203.

Yasugi, T., Umetsu, D., Murakami, S., Sato, M. and Tabata, T. (2008). Drosophila optic lobe neuroblasts triggered by a wave of proneural gene expression that is negatively regulated by JAK/STAT. Development 135, 1471–1480.

Zhou, L. and Luo, H. (2013). Replication protein a links cell cycle progression and the onset of neurogenesis in Drosophila optic lobe development. J Neurosci 33, 2873–2888.

Zhu, S., Lin, S., Kao, C. F., Awasaki, T., Chiang, A. S. and Lee, T. (2006). Gradients of the Drosophila Chinmo BTB-zinc finger protein govern neuronal temporal identity. Cell 127, 409–422.

